# DDAP: docking domain affinity and biosynthetic pathway prediction tool for type I polyketide synthases

**DOI:** 10.1101/637405

**Authors:** Tingyang Li, Ashootosh Tripathi, Fengan Yu, David H. Sherman, Arvind Rao

## Abstract

**Summary:** DDAP is a tool for predicting the biosynthetic pathways of the products of type I modular polyketide synthase (PKS) with the focus on providing a more accurate prediction of the ordering of proteins and substrates in the pathway. In this study, the module docking domain (DD) affinity prediction performance on a hold-out testing data set reached AUC = 0.88; the MRR of pathway prediction reached 0.67. DDAP has advantages compared to previous informatics tools in several aspects: (i) it does not rely on large databases, making it a high efficiency tool, (ii) the predicted DD affinity is represented by a probability (0 to 1), which is more intuitive than raw scores, (iii) its performance is competitive compared to the current popular rule-based algorithm. To the best of our knowledge, DDAP is so far the first machine learning based algorithm for type I PKS pathway prediction. We also established the first database of type I modular PKSs, featuring a comprehensive annotation of available docking domains information in bacterial biosynthetic pathways.

**Availability and implementation:** The DDAP database is available at https://tylii.github.io/ddap. The prediction algorithm DDAP is freely available on GitHub (https://github.com/tylii/ddap) and released under the MIT license.

## 1 Introduction

Natural products (NPs) such as penicillin, erythromycin, artemisinin, taxol and tetrodotoxin are small chemical compounds produced by bacteria, fungi, plants, and animals. These small molecules are found to display a broad range of bioactivities (Katz and Baltz, 2016). About 40% of new drugs approved in the past 30 years were either unaltered NPs or derived from NPs (Newman and Cragg, 2016). Interest is growing in the search for novel NPs in both industrial and academic fields. Although new technologies have improved efficiency, traditional NP discovery requires compound isolation, mass spectrometry (MS) analysis, and nuclear magnetic resonance (NMR) data acquisition, and structure elucidation can be time consuming. Developing computational tools to predict NP structures based on DNA/protein sequences (e.g., antiSMASH (Blin *et al.*, 2017), NP.searcher (Li *et al.*, 2009), PRISM (Skinnider *et al.*, 2017) and SeMPI (Zierep *et al.*, 2017)) is of high interest to facilitate structure characterization, and offer new automated approaches.

Polyketide synthases (PKSs) are one of the most important classes of biosynthetic enzymes. Type I modular PKS (T1PKS) consists of a series of genes encoding multifunctional proteins, including a loading module and multiple extension modules (Dutta *et al.*, 2014). Each extension module is responsible for adding one acyl-monomer to the polyketide chain. The assembly order of polyketide substrates is not always coincident with gene cluster architecture in the bacterial genome. Therefore, finding the correct order of modules and substrates in the polyketide biosynthetic pathway is a crucial step in structure prediction. Previous research has demonstrated that the substrate assembly order is determined by cognate docking domain pairs (DDs) at the N-/C-terminus of PKS proteins (Gokhale and Khosla, 2000; Gokhale *et al.*, 1999). In 2009, Yadav et al. published a rule-based affinity prediction algorithm, based on a general assumption about the 6-deoxyerythronolide B synthase (DEBS) DD structure (Yadav *et al.*, 2009). This method is used by many well-known NP discovery tools including antiSMASH and NP.searcher, despite its several defects (Supplementary Materials). To our knowledge, it is the only available algorithm that specifically predicts DD affinity in T1PKS.

In this study, we collected pathway information of T1PKS from manuscripts published over the past 24 years (1995 - 2018) and developed a machine learning based docking domain affinity prediction tool, DDAP. DDAP uses protein sequences to predict the assembly order of the compounds produced by T1PKS. The DDAP database includes 172 T1PKS and 764 docking domains. As far as we are aware, this is the first and most comprehensive database of DDs in bacterial type I modular PKSs.

## 2 Methods

### 2.1 The DDAP Database

The DDAP database contains 172 records of type I modular PKS proteins, among which 92 are annotated with published pathway information, 80 are annotated with pathways predicted by DDAP. Pathway information includes the docking domain sequences (764 DDs) and the order of genes in the biosynthetic pathway. Users are able to download all above-mentioned datasets from the database and browse the pathway data through an interactive table on the web page.

### 2.2 Docking Domain Affinity Prediction

DDAP has two main functions: (i) predicting the likelihood of interaction given the AA sequences of a C-terminal DD (Head) and a N-terminal DD (Tail); (ii) predicting the most likely pathways based on the predicted DD affinity. DDAP uses machine learning models to predict DD affinity (Fig. S1A, Fig. S1B, Fig. S1C). DDAP takes DD sequences in FASTA format as input and returns the predicted affinity of each DD pair. The affinity will always be a number between 0 and 1, where 1 indicates high affinity (Fig. S1D).

### 2.3 Pathway Prediction

AntiSMASH 4 is a state-of-the-art tool for genome mining for natural product gene cluster discovery (Blin *et al.*, 2017). Following antiSMASH identification of a biosynthetic gene cluster, DDAP can read the PKS annotation and its docking domain affinity prediction algorithm predicts the order of modules/substrates in the biosynthetic pathway (see Supplementary Materials). Alternatively, users can also provide the AA sequences of PKS proteins in FASTA, or CSV format. In the output, DDAP returns an exhaustive list of all possible pathways, each associated with a probability score (0 to 1), and a SMILES string representing the backbone structure of the proposed biosynthetic product. DDAP also provides a plot of the compound structure for the top ten most likely pathways (Fig. S1E).

## 3 Results

According to the five-fold cross-validation results of the best performing model, the area under the receiver operating characteristic (ROC) curve (AUC) for DD affinity prediction was 0.80 (95% CI: 0.78-0.81). The Mean Reciprocal Ranking (MRR) of the true pathways was 0.63 (95% CI: 0.59-0.67). Approximately 71% of the time, the true pathway ranked among the top three. We further tested the best performing model on the hold-out testing set. The final model achieved AUC = 0.88 (95% CI: 0.77-0.98) for DD affinity prediction. The MRR for pathway prediction was 0.67 (95% CI: 0.27-1.00). The true order received the highest likelihood score in 4 of 7 testing pathways (see Supplementary Materials).

Finally, we compared our method with the most widely used method, which was originally developed by Yadav *et al.*, and later adopted by antiSMASH and NP.searcher. We used antiSMASH 4.2.0 to test the performance of Yadav’s method. Seventy pathways were used to compare the performance (see Supplementary Materials). For these 70 pathways, antiSMASH achieved MRR = 0.48 (95% CI: 0.38-0.57); DDAP achieved MRR = 0.62 (95% CI: 0.57-0.66).

## 4 Conclusion

In this study, we established a database for pathways and docking domains of type I modular polyketide synthases. We also built a machine learning based algorithm that predicts T1PKS pathways. The DDAP algorithm is shown to outperform the state-of-the-art without relying on large databases of proteins/compounds (e.g. SeMPI). The prediction tool can be readily incorporated into natural product discovery pipelines and used as a complementary tool along with genome mining software to provide accurate predictions of bacterial type I modular PKS pathways and backbone structures of the secondary metabolites.

