## Supplementary Materials for "DDAP: docking domain affinity and biosynthetic pathway prediction tool for type I polyketide synthases"

### 1 Methods

#### 1.1 Positive and Negative samples

The PKS pathway data used in this study was obtained from two sources: (i) the work reported by Thattai *et al.*, 2007, (ii) a manual literature review, guided by MIBiG database (Version 1.4 and 1.3) (Medema *et al.*, 2015). The PKS sequences were downloaded from the NCBI database. Docking domain sequences were extracted from the N-/C- terminus of the PKS proteins. One hundred amino acids (AAs) at the C-terminus of each PKS protein was extracted as Head (or C-terminal DD) except the last protein in the pathway (which includes the thioesterase (TE) domain). Fifty AAs at the N-terminus of each PKS protein was extracted as the Tail (or N-terminal DD) except the first protein in the pathway (which includes the loading module).

In this study, the ground truth for docking domain (DD) affinity is generated based on the polyketide synthase (PKS) pathway information. Positive samples (encoded as 1) are defined as interacting Head-Tail (HT) pairs (**Figure S1A**). Negative samples (encoded as 0) are defined as Head-Tail pairs that are in the same pathway but not interacting. This notion was also adopted in the study by Thattai et al. (Thattai *et al.*, 2007). Since the total number of negative samples greatly exceeds the total number of positive samples, we randomly subsampled the negative samples to match the size of positive samples to ensure balance.

#### 1.2 Model Selection

During the development stage of DDAP, the dataset was divided into two portions: the training data (85/92 pathways, 714/764 DDs) and the hold-out testing data (7/92 pathways, 50/764 DDs). The training data came from MIBiG Version 1.3 and the Thattai *et al.* dataset (Thattai *et al.*, 2007), which consists of PKS pathways reported before 2016. The hold-out testing data came from MIBiG Version 1.4, which consists of PKS pathways reported between 2016 and 2018. In this study, the names of the PKS pathways were coded as BGC_n, where n was an integer starting from 1. BGC_1 through BGC_85 were used for model selection. BGC_85 through BGC_92 were the hold-out testing samples. Eventually, the best performing model in the CV was trained using all available data (92/92 pathways, 764/764 DDs) and was integrated into DDAP. A mapping scheme from the above IDs to the sequences can be found in the DDAP database.

Model selection is performed through repeated five-fold cross-validation (CV). The area under the receiver operating characteristic (ROC) curve (AUC) is used to assess the DD affinity prediction performance. In CV, the training dataset is randomly partitioned into five approximately equally sized sets, where HT pairs in the same pathway are always allocated to the same set. The five sets are then sequentially used as the validation set, while the remaining four sets are used as the training set. The algorithm learns from the training set and uses the validation set to check the accuracy by calculating the AUC. The above process is repeated five times for each candidate model. In total, $5\ (repeats) \times 5\ (folds) = 25$ AUC scores are calculated. The mean of 25 AUCs is then used to represent the performance of the candidate model. RandomForestRegressor (scikit-learn 0.19.1) is used as the base learner. Parameters are kept as default, except n_estimators (set to 200), random_state (set to 0), and max_depth (a hyperparameter). In the following text, the CV results for the grid search of max_depth is omitted.

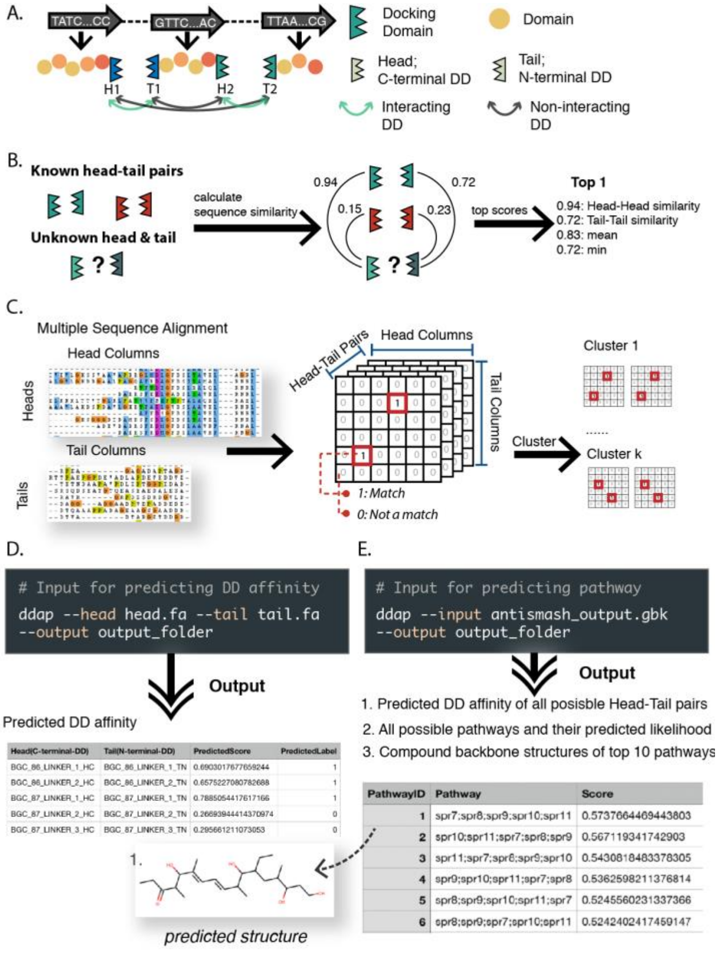


**Figure S1. Overview of DDAP algorithm. A**. A schematic view of polyketide synthase modules and docking domains. **B**. The pipeline of the feature selection method in Model 1. The similarity measure utilized in this study is the bit-score which is not in the range of (0,1) as what is shown in the plot. **C**. The pipeline of the feature selection method in Model 2. **D**. An example of the input and output of DDAP for docking domain affinity prediction. **E**. An example of the input and output of DDAP for pathway prediction.

### 1.3 Model 1: Sequence Similarity Based Model

#### 1.3.1 Method

The underlying assumption of Model 1 is that if an unknown HT pair is similar to an interacting HT pair, the unknown HT pair is then more likely to interact. This assumption is supported by the study of Thattai et al. (Thattai *et al.*, 2007), where it was found that Heads and Tails could be grouped into mutually incompatible clusters based on phylogenetic analysis. For each testing HT pair, Model 1 first calculates the sequence similarity between the unknown Head and all the Heads in the training set, and the similarity between the unknown Tail and all the Tails in the training set (**Figure S1B**). In this study, we use bit-scores calculated by "blastp" in BLAST 2.7.1 to represent sequence similarity. Bit-score is a normalized alignment score that can be used as a measurement for sequence similarity (Altschul *et al.*, 1997). Higher bit-scores represent higher similarity. For each Head-Tail pair in the training set, we further calculate the mean and minimum of the Head-Head similarity and the Tail-Tail similarity. Each interacting HT pair (i.e., each positive sample) in the training set is therefore associated with four scores: (a) the Head-Head similarity score, (b) the Tail-Tail similarity score, (c) the average of (a) and (b), (d) the minimum of (a) and (b).

After obtaining the four metrics, we rank training HT pairs according to (c) in descending order. HT pairs at the top of the list are considered most similar to the unknown (i.e., testing) HT pair. We then use (a), (b), (c) and (d) scores associated with the top $n$ HT pairs as features (**Figure S1B**). Each testing sample will thus be represented by $n \times 4$ numerical features, where $n$ is a hyperparameter optimized through cross-validation. When we calculate features for the training samples, each pathway is treated as a unit, which means that the within-pathway similarity scores are not considered. Considering only inter-pathway similarity ensures that the distribution of features in the training set resembles that of the testing set.

##### 1.3.2 Model 1 CV Results

The Spearman correlation between the four metrics calculated in Model 1 and the ground truth is shown in **Figure S2**. It is observed that the minimum of two similarity scores and the Tail-Tail similarity score have relatively stronger correlation with the ground truth, where positive samples (interacting HT pairs) tend to have higher similarity values. The cross-validation results for Model 1 are shown in **Figure S3**, where we tested $n = 1, 2, 3, 4$. Eventually, $n = 3$ received the highest performance. It is expected that more features receive higher performance. In this case, the improvement from $n = 3$ to $n = 4$ is insignificant, therefore, we consider that the model performance converges at $n = 3$ and $n = 4$.

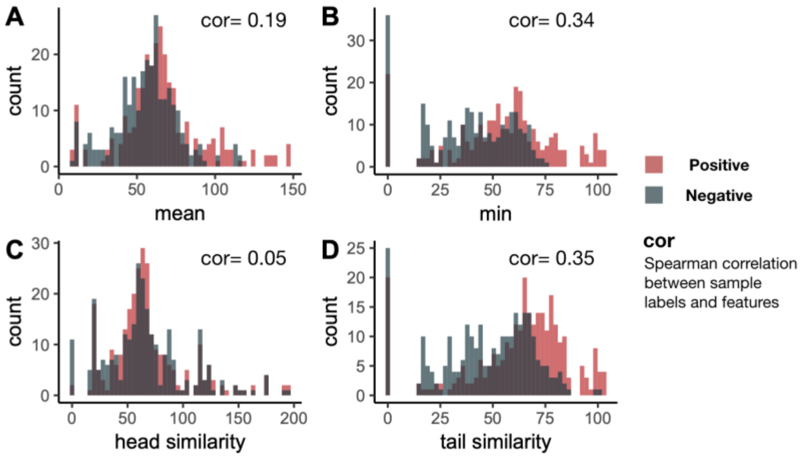


**Figure S2. Histograms of the four metrics calculated in Model 1. A.** Histogram of the mean of the Head-Head similarity and Tail-Tail similarity. **B.** Histogram of the minimum of the Head-Head similarity and Tail-Tail similarity. **C.** Histogram of Head-Head similarity **D.** Histogram of Tail-Tail similarity. "Positive" refers to interacting HT pairs. "Negative" refers to non-interacting HT pairs.

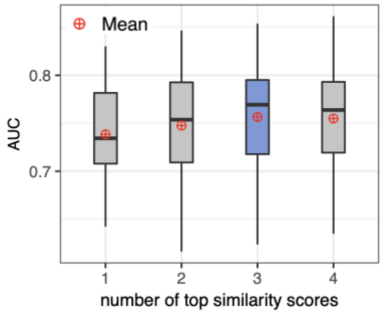


| Top N | AUC |
|---|---|
| 3 | 0.757 |
| 4 | 0.755 |
| 2 | 0.748 |
| 1 | 0.738 |

**Figure S3. A boxplot of the cross-validation results for Model 1.** Each box shows a five-number summary of the cross-validation result of one value of $n$. The AUC in the table is the mean of AUCs in the cross-validation.

### 1.4 Model 2: Cognate Residue Based Model

#### 1.4.1 Method

Previous structural studies on docking domains revealed that key residues (or cognate residues) at the interface of two interacting DD proteins (Broadhurst *et al.*, 2003; Buchholz *et al.*, 2009) are playing important roles in determining the specificity of the interaction and stabilizing the bond. Yadav *et al.* incorporated the above knowledge in their method, assuming the structure of DEBS DD can be generalized to all types of DDs (Yadav *et al.*, 2009). Later on, the crystal structure of type II DD was resolved (Whicher *et al.*, 2013), which was found to have cognate residues at different positions compared to type I DD. On the other hand, Model 2 strives to make use of the knowledge about cognate residues without assuming the locations of the cognate residues. Model 2 contains the following assumptions: (a) Cognate residues in Heads and Tails determine the specificity of Head-Tail interactions; (b) Different classes of DDs have different cognate residues; (c) For an interacting HT pair, correct cognate residues in the Head "match" with cognate residues in the Tail.

In this study, we assume a favorable "match" happens when a positively charged amino acid (AA) meets a negatively charged AA. As shown in **Figure S1C**, matched and unmatched residue pairs can be represented by 1 or 0 in a 2-D matrix, where "1" marks the locations of matched residue pairs. For example, "1" at location $[2,5]$ means the second AA in the Head matches with the fifth AA in the Tail. After generating the 2-D representation of each pair of DDs, we use k-means to cluster these 2-D matrices into k clusters (k is a hyperparameter) to address assumption (b). The 2-D matrices are turned into 1-D vectors before the k-means clustering.

After clustering, HT pairs that have different locations of cognate residues will be allocated to different clusters. In order to find cognate residues within each cluster, we rank each location $[i, j]$ in the 2-D matrix by the frequency of observing a "1" at that position, which can be calculated as:

$$p_{i,j} = \frac{\sum_{n=1}^{N} I_{match}(i,j,n)}{N}$$

$$I_{match}(i,j,n) = \begin{cases} 1 \; if \; the \; i^{th} \; Head \; residue \; matches \; with \; the \; j^{th} \; Tail \; residue \; for \; the \; N^{th} \; DD \; pair \\ 0 \qquad\qquad\qquad\qquad\qquad\qquad\qquad\qquad\qquad\qquad\qquad\qquad otherwise \end{cases}$$, where

N is the total number of HT pairs. The locations with the highest frequencies are picked as cognate residue pairs for that particular cluster. The total number of cognate residues in a cluster is a hyperparameter. We used a combination of cognate residue pairs from all clusters as the features for Model 2. Repeated cognate residue pairs are removed.

#### 1.4.2 Model 2 CV Results

We used principal component analysis (PCA) to visualize the clustering result in 2-D and 3-D space (**Figure S4**). Although our approach is different from the phylogenetic approach that Thattai et al. utilized and we are using a different dataset, we still observed that the docking domain pairs forms three large clusters. **Figure S5** showed the cross-validation result for the grid search of the number of HT pair clusters ($k$) and the number of cognate residues in each cluster ($c$). As the two parameters increase, the mean AUC slowly converges to 0.72. We picked $k = 7$ and $c = 29$ as the parameters for the final model. Aside from k-means, we also tested DBSCAN, k-modes and spectral clustering. These three methods did not outperform the proposed method (results not shown).

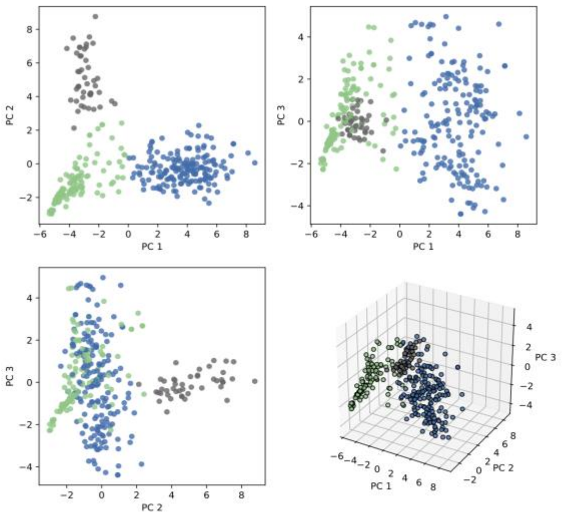


**Figure S4. Clustering result for k = 3.** Principal component analysis (PCA) is used to generate the 2-D and 3-D visualization of the clustering result. In this figure, each dot represents a HT pair. The data used as the input for PCA represents the features generated by Model 2. The color of the dots indicates cluster membership.

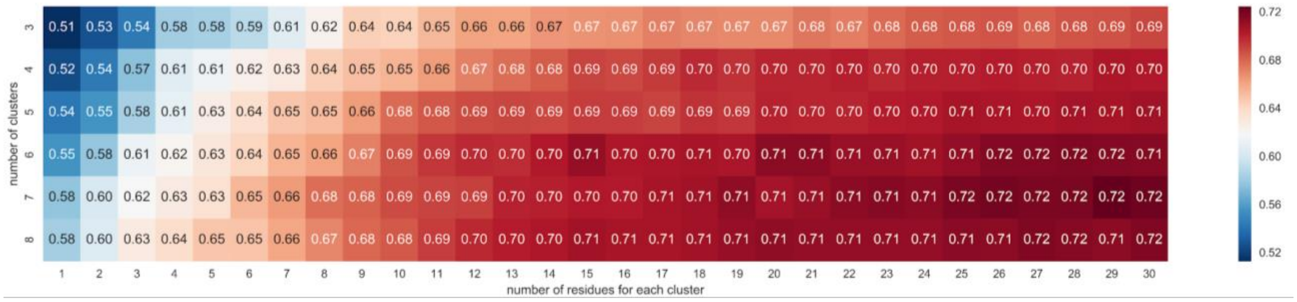


**Figure S5. Cross-validation results for the grid search of hyperparameters in Model 2.** The value and color in each grid cell represent the average AUC for a specific hyperparameter combination. The vertical axis is the number of clusters. The horizontal axis is the number of cognate residues picked for each cluster.

We then stacked the prediction results of Model 1 and Model 2, in order to reduce biases and boost the prediction accuracy. As stated above, Model 1 and Model 2 are two independent DD affinity prediction models. Each testing sample (i.e. each unknown HT pair) will receive two predicted affinity scores from the two models. To combine the two scores, we used weighted averaging, which is a widely used ensemble method for regression models. The weights are tuned through CV (**Figure S6**). The highest AUC (0.7977) was achieved when the weight of Model 2 $w_2 = 0.6$ and the weight of Model 1 $w_1 = 0.4$. Aside from weighted averaging, Support Vector Machines (SVM) and Random Forest (RF) were employed as alternative ensemble methods (results not shown). We also tried combining features generated by Model 1 and Model 2 into one feature matrix and train RF with the entire matrix (results not shown). Weighted averaging performed best among all candidate methods and thus was used in the final model.

#### 1.5 Pathway Prediction

We then used the predicted DD affinity to infer PKS protein assembly orders in the biosynthetic pathway. For a given PKS gene cluster, we first create an exhaustive list of putative pathways generating permutations of PKS proteins. Then for each candidate pathway we calculate average affinity of all interacting HT pairs in the pathway: $\frac{\sum_{i=1}^{N-1} f_a(H_i,T_{i+1})}{N-1}$, where function $f_a(H,T)$ calculates the affinity between $H$ and $T$; $N$ is the total number of proteins in the pathway; $H_i$ is the C-terminal DD of the $i$-th protein in the putative pathway; $T_i$ is the N-terminal DD of the $i$-th protein in the putative pathway. The likelihood score associated with each pathway always ranges from 0 to 1, where 1 indicates high likelihood. Eventually, we rank the putative pathways according to the likelihood. Since a brute-force approach is used for pathway prediction, the running time grows exponentially with the number of genes/proteins in the biosynthetic pathway. Because of limited time and resources, in cross-validation we only predicted pathways for PKSs with ten or less genes. BGC_18, BGC_27 and BGC_29 were omitted because they have more than ten genes. Pathway prediction performance in the CV of the final model is shown in a pie chart in **Figure S7**. The Mean Reciprocal Rank (MRR) of the true pathway is 0.63 (95% CI: 0.59-0.67). Pathway prediction performance on the hold-out testing dataset is shown in **Table S1**.

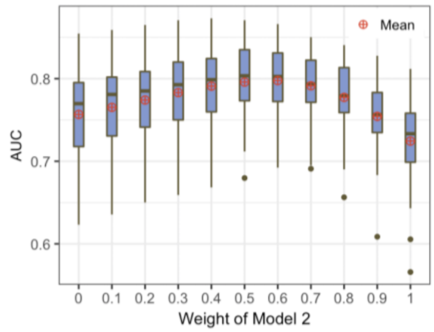


| Weight | AUC |
|---|---|
| **0** | 0.7567 |
| **0.1** | 0.7652 |
| **0.2** | 0.7741 |
| **0.3** | 0.7831 |
| **0.4** | 0.7910 |
| **0.5** | 0.7959 |
| **0.6** | 0.7977 |
| **0.7** | 0.7914 |
| **0.8** | 0.7773 |
| **0.9** | 0.7544 |
| **1** | 0.7244 |

**Figure S6. Cross-validation results for the stacked model.** This figure summarizes CV results when the weight of Model 2 ($w_2 \in [0,1]$) takes different values. The weight of Model 1 is $w_1 = 1 - w_2$. In this plot, $w_2 = 1$ is equivalent to using only Model 2; $w_2 = 0$ is equivalent to using only Model 1. Among all weights, $w_2 = 0.6$ achieved the highest average AUC and was used in the final model.

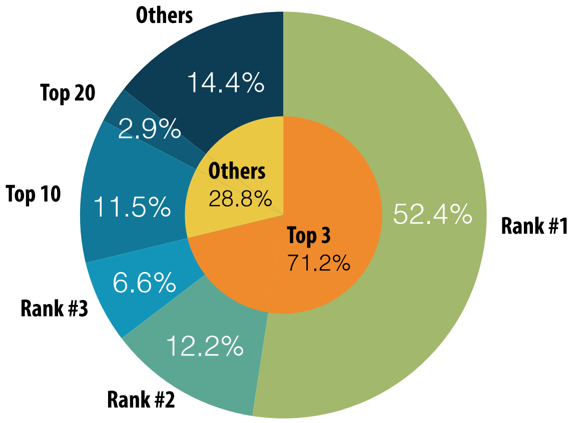


**Figure S7. Cross-validation results of the final model for pathway prediction.** This figure shows the frequency for the true pathway ranking #1, #2, etc., in our ranking of putative pathways. In the outer pie chart, the "Top 10" category includes #4 through #10. "Top 20" includes #11 through #20. "Others" refers to the cases where the ranks of true pathways are beyond #20. In the inner pie chart, "Top 3" includes #1 through #3; "Others" refers to ranks beyond #3.

#### 1.6 Comparing with the State-of-the-art

The rule-based method that Yadav *et al.* developed is the current state-of-the art for pathway prediction for T1PKS (Yadav *et al.*, 2009). However it has several defects: (i) This algorithm assumes that the structure of DEBS DD can be applied to other DDs, however, it has been proven that the DD structure of DEBS does not apply to all DDs (Whicher *et al.*, 2013), (ii) The algorithm typically yields a large number of tied ranks in pathway prediction. (iii) The upper/lower limits of the pathway scores calculated by this algorithm are different for PKSs with different numbers of modules/genes, which makes the scores difficult to interpret. DDAP has overcome all the above limitations.

We used antiSMASH 4.2.0 web server (Blin *et al.*, 2017) for the pathway prediction performance comparison. By default, for each PKS system with more than three and less than 11 PKS genes, antiSMASH returns a maximum of 1000 putative pathways and the raw scores associated with each putative pathway. For PKSs with $\leq 3$ or $\geq 11$ genes, collinearity will be assumed, meaning that the order of PKS proteins in the pathway is assumed to be the same as the order of their coding genes in the genome. AntiSMASH will not predict putative PKS pathways when non-PKS genes (e.g. NRPS) are present in the gene cluster. Therefore, we removed all non-PKS genes before uploading the sequences to antiSMASH server to ensure that the pathway prediction was performed.

We first attempted to use the hold-out testing data to compare the performance (see **Table S1**), and then calculated the Mean Reciprocal Rank (MRR) for antiSMASH and our algorithm. The antiSMASH method typically introduces tied scores to the results. Therefore, we used a tie-aware MRR to measure the performance of antiSMASH (McSherry and Najork, 2008). A conventional MRR was used for the evaluation of our method, which is equivalent to using the tie-aware MRR with the number of tied scores set to 1. For the hold-out testing data, MRR of the antiSMASH method was 0.64; MRR of our method was 0.67.

| BGC ID | rank of true pathways (DDAP) | rank of true pathways (antiSMASH) | tied pathways (antiSMASH) | predicted or assumed colinear (antiSMASH) | total number of possible pathways | number of genes in the pathway |
|---|---|---|---|---|---|---|
| **BGC_86** | 1 | 1 | 1 | assumed colinear | 6 | 3 |
| **BGC_87** | 1 | 1 | 2 | predicted | 24 | 4 |
| **BGC_88** | 26 | 1000+ | / | predicted | 40320 | 8 |
| **BGC_89** | 1 | 1 | 2 | predicted | 120 | 5 |
| **BGC_90** | 1 | 1 | 1 | assumed colinear | 6 | 3 |
| **BGC_91** | 6 | 120+ | / | predicted | 720 | 6 |
| **BGC_92** | 2 | 1 | 1 | assumed colinear | 6 | 3 |

**Table S1. Pathway prediction results of DDAP and antiSMASH on hold-out testing dataset.** The relationship between the total number of possible pathways ($P$) and the number of genes in the pathway ($g$) is $P = g!$. In the third column, $x+$ means that $x$ possible pathways are returned by antiSMASH, but the true pathway is not among these putative pathways. When $x < 1000$, it could be because: (i) the predicted PKS genes are incorrect, (ii) antiSMASH uses a separate method to locate the first and last module before generating permutations of the PKS proteins. If the first/last module is incorrectly predicted, the result will consist of a non-exhaustive list of putative pathways that do not include the true pathway. The number of tied pathways (pathways that have the exact same score as the true pathway) is shown in the fourth column.

Since the hold-out testing set is limited in size for a robust performance comparison, we further used training pathways to compare antiSMASH and DDAP. The antiSMASH web server was used to make predictions for the training pathways (i.e., the pathways used for CV of our method). We excluded 15 samples (see reasons in **Table**

**S2**). Eventually, 70 samples were used for the performance comparison. To calculate MRR for DDAP on these 70 samples, we extracted all predictions made for these samples in the cross-validation of our best performing model. Eventually, our method achieved MRR=0.62 (95% CI: 0.57-0.66), while the antiSMASH method achieved MRR=0.48 (95% CI: 0.38-0.57). Since PKSs with assumed colinearity do not directly reflect the performance of the prediction algorithm, we calculated MRR a second time after removing the PKSs that were assumed to be colinear (**Table S2**). Eventually, antiSMASH received MRR=0.31 (95% CI: 0.23-0.40), and our method received MRR=0.54 (95% CI: 0.49-0.59) (**Figure S8**). Note that during CV, the DDAP algorithm was trained using 80% of the training data (about 74% of all available data). This result shows that even when DDAP is trained with 74% of data, it still outperforms antiSMASH in most cases.

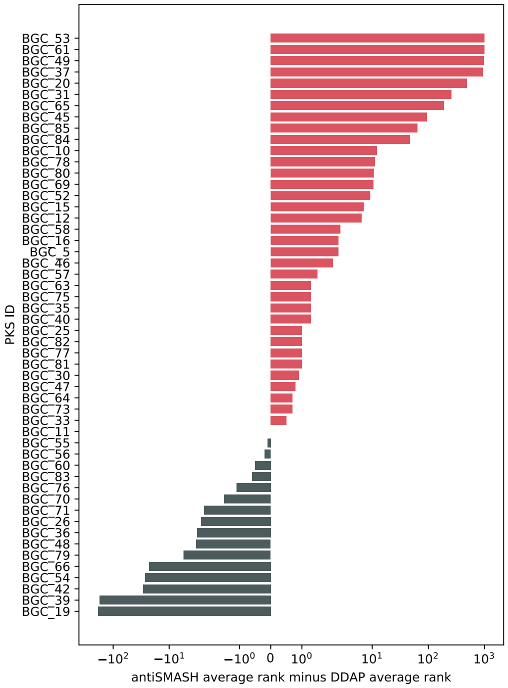


**Figure S8. Pathway prediction results of DDAP and antiSMASH on CV dataset.** The red bars represent the cases where DDAP outperforms antiSMASH. The average rank for antiSMASH is calculated as: $r_h + \frac{t-1}{2}$, where $t$ is the number of tied pathways ($t \geq 1$); $r_h$ is the highest possible rank of the true pathway. The average rank for DDAP is simply the average of the predicted ranks in the CV results. All PKSs with assumed colinearity are removed from this plot. BGC_13 is not included in this plot because the average rank predicted by antiSMASH and DDAP both exceed 1000.

### 1.7 Alternative Methods

A commonly used approach for protein-protein interaction (PPI) prediction is coevolution analysis (de Juan *et al.*, 2013). RaptorX (Källberg *et al.*, 2012) is one of the most acknowledged tools in this field. The "Complex Contact Prediction" function in RaptorX uses coevolution analysis and deep learning to predict the interfacial contacts between two potentially interacting proteins. We used the RaptorX webserver to obtain contact maps of both positive and negative samples (i.e., interacting and non-interacting DDs) in the training set, and used deep neural networks implemented by Google AutoML Vision to predict DD affinity based on the contact maps. We found this method achieved an accuracy of around 50%. Therefore, this approach was not included in the final model.

| BGC ID | rank of true pathways | tied pathways | include or exclude for performance comparison | notes |
|---|---|---|---|---|
| **BGC_14** | 1 | 1 | include | assumed colinear |
| **BGC_17** | 1 | 1 | include | assumed colinear |

| | | | | |
|---|---|---|---|---|
| **BGC_23** | 1 | 1 | include | assumed colinear |
| **BGC_24** | 1 | 1 | include | assumed colinear |
| **BGC_28** | 1 | 1 | include | assumed colinear |
| **BGC_38** | 1 | 1 | include | assumed colinear |
| **BGC_4** | 1 | 1 | include | assumed colinear |
| **BGC_41** | 1 | 1 | include | assumed colinear |
| **BGC_44** | 1 | 1 | include | assumed colinear |
| **BGC_50** | 1 | 1 | include | assumed colinear |
| **BGC_59** | 1 | 1 | include | assumed colinear |
| **BGC_6** | 1 | 1 | include | assumed colinear |
| **BGC_62** | 1 | 1 | include | assumed colinear |
| **BGC_67** | 1 | 1 | include | assumed colinear |
| **BGC_68** | 1 | 1 | include | assumed colinear |
| **BGC_7** | 1 | 1 | include | assumed colinear |
| **BGC_74** | 1 | 1 | include | assumed colinear |
| **BGC_58** | 7 | 12 | include | predicted |
| **BGC_10** | 3 | 22 | include | predicted |
| **BGC_11** | 1 | 1 | include | predicted |
| **BGC_12** | 3 | 10 | include | predicted |
| **BGC_13** | 1000+ | / | include | predicted |
| **BGC_15** | 5 | 10 | include | predicted |
| **BGC_16** | 3 | 2 | include | predicted |
| **BGC_19** | 1 | 12 | include | predicted |
| **BGC_20** | 871 | 134 | include | predicted |
| **BGC_25** | 2 | 1 | include | predicted |
| **BGC_26** | 5 | 6 | include | predicted |
| **BGC_30** | 1 | 4 | include | predicted |
| **BGC_31** | 49 | 624 | include | predicted |
| **BGC_33** | 1 | 2 | include | predicted |
| **BGC_35** | 1 | 4 | include | predicted |
| **BGC_36** | 1 | 1 | include | predicted |
| **BGC_37** | 1000+ | / | include | predicted |
| **BGC_39** | 275 | 526 | include | predicted |
| **BGC_40** | 1 | 6 | include | predicted |
| **BGC_42** | 17 | 8 | include | predicted |
| **BGC_45** | 63 | 148 | include | predicted |
| **BGC_46** | 10 | 5 | include | predicted |
| **BGC_47** | 2 | 1 | include | predicted |
| **BGC_48** | 1 | 2 | include | predicted |
| **BGC_49** | 2+ | / | include | predicted |
| **BGC_5** | 1 | 6 | include | predicted |
| **BGC_52** | 10 | 19 | include | predicted |
| **BGC_53** | 6+ | / | include | predicted |
| **BGC_54** | 25 | 102 | include | predicted |
| **BGC_55** | 1 | 2 | include | predicted |

| | | | | |
|---|---|---|---|---|
| **BGC_56** | 2 | 3 | include | predicted |
| **BGC_57** | 1 | 4 | include | predicted |
| **BGC_60** | 1 | 2 | include | predicted |
| **BGC_61** | 24+ | / | include | predicted |
| **BGC_63** | 1 | 4 | include | predicted |
| **BGC_64** | 1 | 4 | include | predicted |
| **BGC_65** | 169 | 168 | include | predicted |
| **BGC_66** | 4 | 9 | include | predicted |
| **BGC_69** | 9 | 6 | include | predicted |
| **BGC_70** | 1 | 2 | include | predicted |
| **BGC_71** | 2 | 1 | include | predicted |
| **BGC_73** | 1 | 4 | include | predicted |
| **BGC_75** | 1 | 4 | include | predicted |
| **BGC_76** | 1 | 6 | include | predicted |
| **BGC_77** | 2 | 3 | include | predicted |
| **BGC_78** | 13 | 4 | include | predicted |
| **BGC_79** | 5 | 2 | include | predicted |
| **BGC_80** | 5 | 16 | include | predicted |
| **BGC_81** | 1 | 3 | include | predicted |
| **BGC_82** | 7 | 11 | include | predicted |
| **BGC_83** | 1 | 1 | include | predicted |
| **BGC_84** | 51 | 31 | include | predicted |
| **BGC_85** | 115 | 6 | include | predicted |
| **BGC_2** | / | / | exclude | hybrid |
| **BGC_21** | / | / | exclude | hybrid |
| **BGC_22** | / | / | exclude | hybrid |
| **BGC_3** | / | / | exclude | hybrid |
| **BGC_32** | / | / | exclude | hybrid |
| **BGC_43** | / | / | exclude | hybrid |
| **BGC_8** | / | / | exclude | hybrid |
| **BGC_34** | / | / | exclude | hybrid |
| **BGC_18** | / | / | exclude | no results from CV |
| **BGC_27** | / | / | exclude | no results from CV |
| **BGC_29** | / | / | exclude | no results from CV |
| **BGC_1** | / | / | exclude | wrong region |
| **BGC_51** | / | / | exclude | wrong region |
| **BGC_72** | / | / | exclude | wrong region |
| **BGC_9** | / | / | exclude | wrong region |

**Table S2. AntiSMASH pathway prediction results.** In the second column, $x+$ means that $x$ possible pathways are returned by antiSMASH, and the true pathway is not among these putative pathways. The third column is the number of pathways tied with the true pathway (including the true pathway itself). The fourth column indicates whether the PKS is considered for performance comparison. In the fifth column, “predicted” means that the pathway is predicted by antiSMASH; “assumed colinear” means the pathway is generated assuming colinearity; “hybrid” means antiSMASH did not make pathway prediction for this gene cluster because hybrid genes are present; “wrong region” means that the PKS regions predicted by antiSMASH are inaccurate; “no results from CV” means that these PKSs did not receive pathway predictions in our CV.
